# Metabolic heritage mapping: heterogenous pools of cytoplasmic nucleotide sugars are selectively utilized by various glycosyltransferases

**DOI:** 10.1101/2021.11.03.467160

**Authors:** Paulina Sosicka, Bobby G. Ng, Lauren E. Pepi, Asif Shajahan, Maurice Wong, David A. Scott, Kenjiroo Matsumoto, Zhi-Jie Xia, Carlito B. Lebrilla, Robert S. Haltiwanger, Parastoo Azadi, Hudson H. Freeze

**Affiliations:** Human Genetics Program, Sanford Burnham Prebys, La Jolla, California; Complex Carbohydrate Research Center, University of Georgia, Athens, Georgia; Department of Chemistry, University of California Davis, Davis, California; Cancer Center, Sanford Burnham Prebys, La Jolla, California

**Keywords:** exogenous substrate, salvage pathway, *de novo* pathway, glycosylation, nucleotide sugars

## Abstract

Biosynthesis of macromolecules requires precursors such as sugars or amino acids, originating from exogenous/dietary sources, reutilization/salvage of degraded molecules or *de novo* synthesis. Since these sources are assumed to contribute to one homogenous pool, their individual contributions are often overlooked. Protein glycosylation uses monosaccharides from all the above sources to produce nucleotide sugars required to assemble hundreds of distinct glycans. Here we demonstrate that cells identify the origin/heritage of the monosaccharide, fucose, for glycosylation. We measured the contribution of GDP-fucose from each of these sources for glycan synthesis and found that different fucosyltransferases, individual glycoproteins, and linkage-specific fucose residues identify and select different GDP-fucose pools dependent on their heritage. This supports the hypothesis that GDP-fucose exists in multiple, distinct pools, not as a single homogenous pool. The selection is tightly regulated since the overall pool size remains constant. We present novel perspectives on monosaccharide metabolism, which may have general applicability.

## INTRODUCTION

Biosynthesis of macromolecules requires precursors such as sugars or amino acids, which are derived alternatively from the diet (exogenous precursors), *via* reutilization (salvage) of degraded molecules or by *de novo* synthesis. Often, salvaged and exogenous precursors are thought to intermingle and are treated a homogenous source, but it is important to emphasize that these two pathways must be considered separately. Exogenous molecules originate from the diet and enter the cell via specific transporter(s) as single molecules, while salvaged precursors are recycled from macromolecules following their lysosomal or proteasomal degradation. The common assumption is that the heritage of these precursors is of little importance since they contribute to a common, homogenous pool intended for different cellular processes.

Monosaccharides are used by cells to produce energy and for synthesis of macromolecules. Glycosylation is the process by which carbohydrates are covalently attached to a target macromolecule utilizing monosaccharides from varying sources to produce nucleotide sugars; substrates that are further used to assemble distinct glycans (Varki et al., 2015). Under physiological concentrations, detailed analysis of mannose metabolism in human fibroblasts demonstrated that neither gluconeogenesis, glycogen, nor mannose salvaged from glycoprotein degradation contributed to mannose found in N-glycans. In contrast, cells efficiently use exogenous mannose for N-glycan biosynthesis (Ichikawa et al., 2014). How cells determine which portion of a monosaccharide pool is directly incorporated into glycans and which fraction is utilized for other cellular processes, i.e., as an energy source, remains unknown. Although efforts have been made to study how these different pathways contribute to nucleotide sugar biosynthesis, they were never investigated simultaneously. Does glycosylation recognize the heritage of the monosaccharide? If yes, how is this regulated? We believe these questions are significant because dietary monosaccharide supplements are successful therapies for some disorders of glycosylation (Verheijen et al., 2020) and even for cancer (Gonzalez et al., 2018).

In humans, fucose is incorporated into glycans using GDP-fucose as a nucleotide sugar donor (Schneider et al., 2017). It can be synthesized directly from exogenous fucose (fucose^Ex^) or from fucose salvaged from glycoprotein degradation (fucose^Sal^) (Coffey et al., 1964; Kaufman and Ginsburg, 1968). Both require subsequent actions of fucokinase (FCSK) and fucose-1-phosphate guanylytransferase (FPGT) to “activate” fucose (Ishihara et al., 1968). Alternatively, GDP-fucose can be produced *de novo* from either mannose (fucose^Man^) or glucose (fucose^Glc^). In this process, GDP-mannose is converted to GDP-fucose in a three-step reaction catalyzed by two enzymes – GDP-mannose 4,6-dehydratase (GMDS) and GDP-fucose synthetase (GFUS) (Ginsburg, 1960) (**Figure 1A**). In 1975, isotope-dilution studies using ^3^H-fucose at a single concentration of 0.3 μM, demonstrated only ~10 % of GDP-fucose in HeLa cells originated from fucose^Ex^, while ~90 % was assumed to be synthesized *de novo* (Yurchenco and Atkinson, 1975, 1977). This method assumed that the radiolabeled component is a metabolic tracer, making little contribution to the overall products. The contribution of fucose^Sal^ (or other monosaccharides) was never addressed.

**Figure 1.**
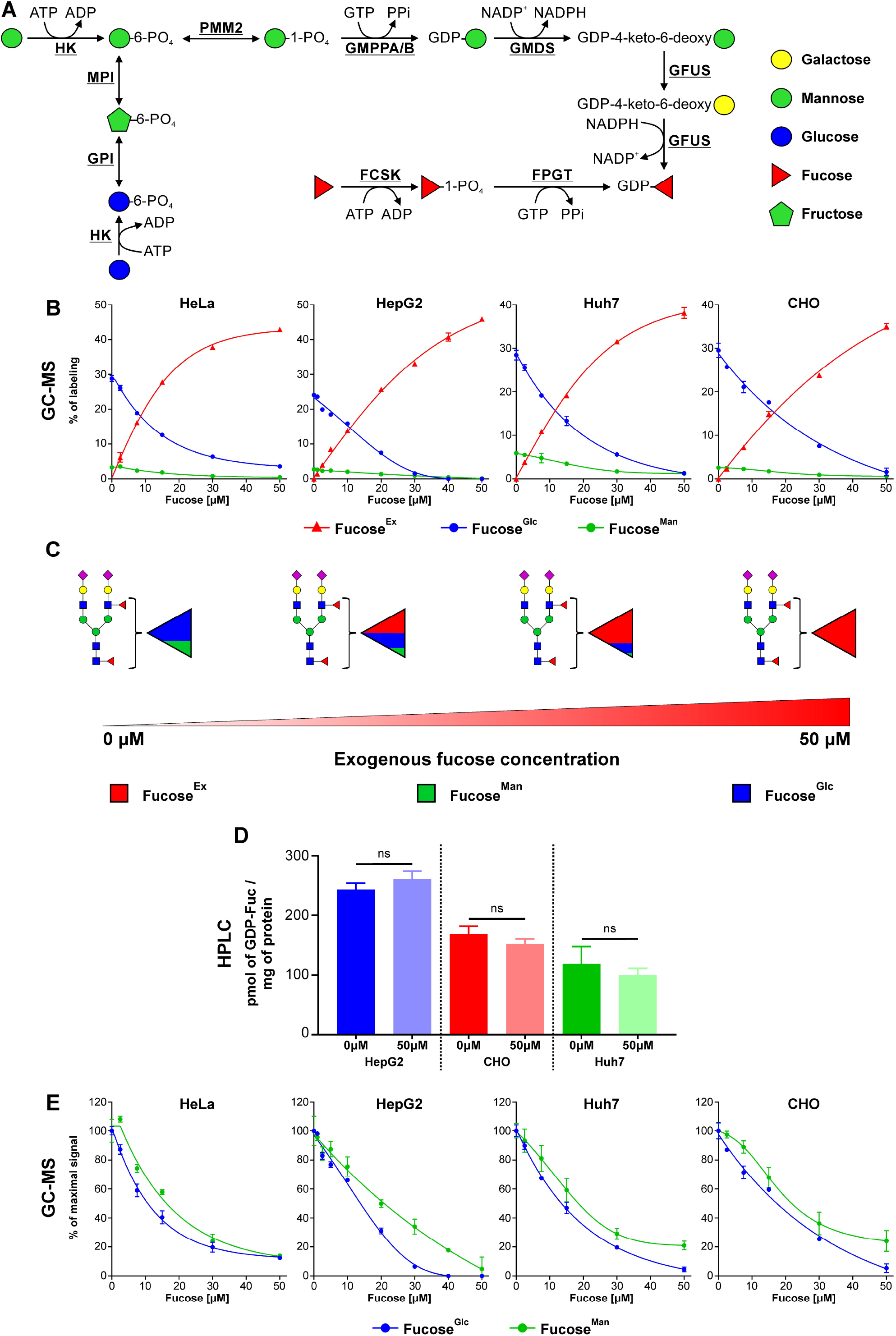
Cells selectively utilize fucose derived from mannose and glucose as well as fucose originating from an exogenous source. (A) Schematic showing the biosynthetic pathways involved in GDP-fucose production. (B and E) Incorporation of 5 mM ^13^C-UL-glucose, 50 μM ^13^C-3,4-mannose and ^13^C-6-fucose into fucosylated N-glycans produced by HeLa, HepG2, Huh7 and CHO cells expressed as a percentage of labeling (B) or as a percentage of maximal signal (E); n = 3; data are presented as mean ± SD. (C) Schematic representation of the results presented in panel B. (D) Comparison of GDP-fucose amount produced by HepG2, CHO and Huh7 treated and untreated with exogenous fucose. Statistically significant was assigned using *t*-test; ns – nonsignificant; ns p > 0.05; n = 5; data are presented as mean ± SD.

GDP-fucose incorporation into glycoconjugates requires its delivery into the Golgi apparatus and endoplasmic reticulum (ER). This process is mediated by two nucleotide sugar transporters, one of them is the major provider of the Golgi GDP-fucose, while the other delivers this nucleotide sugar to the ER (Lu et al., 2010; Lühn et al., 2001). In humans, there are 13 known fucosyltransferases (FUTs), which mostly localize to the Golgi, where they are engaged in fucosylation of N- and O-glycans as well as glycolipids (Schneider et al., 2017). Although many of them have overlapping substrate specificities there is only one enzyme, FUT8, that adds α1-6 fucose to N-glycan chitobiose core (Yang and Wang, 2016). There are two known FUTs localized to the ER, where they add O-fucose directly to Ser/Thr residues of distinct glycoproteins (Schneider et al., 2017).

Using fucose as a model monosaccharide, we evaluated the simultaneous contributions of exogenous, salvaged and *de novo* produced fucose to glycosylation. Our results show cells can tightly regulate multiple pools of GDP-fucose and can recognize their heritage for glycan synthesis.

## RESULTS

### Cells selectively utilize fucose from different sources

We measured the relative contribution of fucose^Ex^ and fucose from the *de novo* pathway by incubating cells with differentially labeled ^13^C-monosaccharides. N-glycan-associated fucose synthesized from exogenous glucose (5 mM), mannose (50 μM) and directly from fucose (0-50μM) was determined following acid hydrolysis and analysis by GC-MS (**Supplementary Figure 1A, 1B, 1C**). Their contributions are assumed to reflect their abundance in the GDP-fucose pool, since it is the only known source for glycan synthesis. Using this GC-MS method, we repeated the HeLa cell experiments (Yurchenco and Atkinson, 1975, 1977) and confirmed a dominant role of the *de novo* pathway at very low fucose concentration ≤2.5 μM. In the absence of fucose^Ex^ ~5 times more N-glycan-associated fucose originates from ^13^C-glucose than from ^13^C-mannose (**Figure 1B, 1C, Supplementary Figure 1D**). Since the physiological concentration of mannose (50 μM) is ~100 times less than glucose (5 mM), mannose is utilized more efficiently for fucosylation compared to glucose, as previously seen for mannose in N-glycans (Ichikawa et al., 2014). However, GDP-fucose produced by the *de novo* pathway is completely suppressed by fucose^Ex^, which becomes the major source of N-glycan-associated fucose, when it is added to ~30-50 μM (**Figure 1B, 1C**). The same pattern was seen in nine different cell lines of varying tissue origin (**Figure 1B, 1C, Supplementary Figure 1D**). To eliminate possible kinetic isotope effects, we exchanged the number and location of ^13^C carbons between glucose, mannose and fucose but the results remained unchanged (**Figure 1B, Supplementary Figure 1F**). Labeling time (12 h and 48 h) did not affect our results either (**Figure 1B, Supplementary Figure 1H**).

The preference of fucose^Ex^ might be explained by an increase in the total GDP-fucose pool. Therefore, we isolated nucleotide sugars from HepG2, Huh7 and CHO cells grown in the absence or presence of 50 μM fucose and demonstrated that the treatment does not increase the total cytosolic concentration of GDP-fucose (**Figure 1D, Supplementary Figure 2**). Therefore, it appears that GDP-fucose originating from fucose^Ex^ suppresses *de novo* GDP-fucose synthesis. The most likely explanation is the known feedback inhibition of the *de novo* pathway enzyme GMDS (**Figure 1A**) by GDP-fucose as previously shown for human and *E. coli* enzymes (Kornfeld and Ginsburg, 1966; Sullivan et al., 1998).

The ratio of glucose to mannose contributions into N-glycan-associated mannose varies from 5: 1 to 3:1 between different cell lines (Ichikawa et al., 2014). This ratio does not change upon fucose^Ex^ titration (**Supplementary Figure 3**). Therefore, we expected that this would also be true for their contribution to N-glycan-associated fucose. However, the contribution of fucose^Glc^ preferentially decreases compared to that from fucose^Man^. Between 0-30 μM fucose^Ex^ the ratio changes from approximately 6:1 to 3:1 in HepG2 cell (**Figure 1E, Supplementary Figure 1E, 1G, 1I**). This is not compatible with a single homogenous GDP-mannose pool serving as a substrate for GMDS. Therefore, we believe there are at least two GDP-mannose pools.

### Exogenous fucose has little impact on the use of salvaged fucose

To measure fucose salvage from degraded glycans, we focused on newly synthesized, secreted, fucosylated N-glycoproteins. This approach allowed us to simultaneously study all four sources of N-glycan-associated fucose: fucose^Man^, fucose^Glc^, fucose^Ex^ and fucose^Sal^. To distinguish fucose^Sal^ from fucose^Ex^, we first pre-labeled cells for 11 days with ^13^C-UL-Fucose (M+6), that replaced over 90 % of ^12^C-fucose, before labeling with exogenous ^13^C-6-fucose (M+1).

To ensure that the secreted material predominantly contains newly synthesized proteins rather than those released by surface proteolysis, we pulse labeled HepG2 cells for 2 h with S^35^ Met to assess total proteins and fucosylated glycoproteins (**Supplementary Figure 4A, 4B**). To prove that secreted radioactivity comes from newly synthesized proteins and not from the material detaching from cells we used brefeldin A (BFA), an inhibitor of the vesicle membrane trafficking (Helms and Rothman, 1992). Treatment with BFA almost completely prevented the secretion of fucosylated proteins confirming applicability of the designed approach (**Supplementary Figure 4B**). In addition, we demonstrated that after approximately 2 h all the labeled material is secreted (**Supplementary Figure 4B**). This was further confirmed by analyzing the secretion of glycoproteins containing ^3^H-fucose after 2 h pulse labeling of HepG2 cells, where all the label material is secreted within ~90 min (**Supplementary Figure 4C**).

We used HepG2, Huh7 and CHO cells pre-labeled with ^13^C-UL-Fucose to measure the contributions of each source of fucose to secreted, fucosylated N-glycans. For both HepG2 and Huh7, the contributions of salvage and *de novo* pathways are comparable in the absence of fucose^Ex^. In contrast, CHO cells rely more heavily on the salvage pathway. As previously observed, fucose^Ex^ quickly suppresses the *de novo* pathway. However, utilization of fucose^Sal^ is much less sensitive to fucose^Ex^ (**Figure 2A, 2B**).

**Figure 2.**
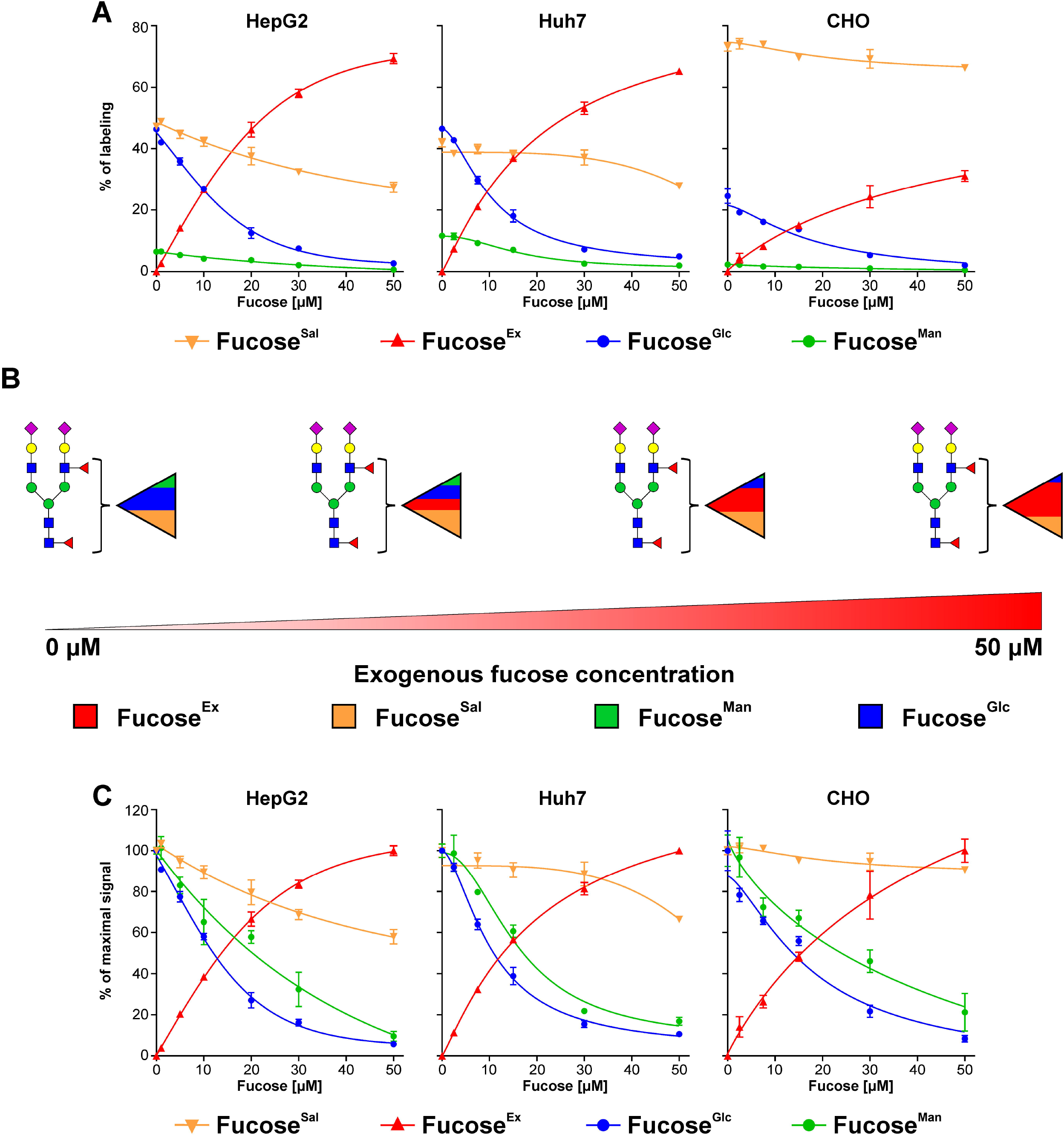
The contribution of fucose salvage into N-glycans is almost insensitive to exogenous fucose. (A and C) Incorporation of 5 mM ^13^C-1,2-glucose, 50 μM ^13^C-1,2,3-mannose and ^13^C-6-fucose into fucose associated with newly synthesized N-glycoproteins secreted by HepG2, Huh7 and CHO cells pre-labeled for eleven days with 50 μM ^13^C-UL-fucose expressed as a percentage of labeling (A) and as a percentage of maximal signal (C); n = 3; data are presented as mean ± SD. (B) Schematic representation of the results presented in panel A.

We found that fucose^Man^ is, once again, less sensitive to fucose^Ex^ than fucose^Glc^. Fucose^Sal^ is treated differently; its contribution changes very little i.e., decreases only by 10-40 % when exposed to fucose^Ex^ (**Figure 2C**). This confirms that cells can distinguish and preferentially incorporate GDP-fucose of one heritage over another. This is compatible with the existence of separate GDP-fucose pools in the cytoplasm.

### Fucose linkage drives the selectivity between different GDP-fucose pools

We investigated the utilization of different GDP-fucose sources in N-glycosylation by first analyzing the contribution of fucose^Ex^ into N-glycans with one and two fucose residues. In N-glycans with a single fucose, this monosaccharide is predominantly attached to the chitobiose core as α1-6 fucose and is referred to as core fucosylation. Only one fucose residue can be attached to the chitobiose core; all others must reside on N-glycan antennae, linked to N-acetylglucosamine via α1-3 or α1-4 linkage or to galactose via α1-2 linkage.

First, we compared the incorporation efficiency of exogenous ^13^C-UL-fucose to ^12^C-fucose that originated from endogenous sources i.e., salvage or *de novo* pathway (fucose^Endo^) using NanoHPLC Chip-Q-TOF MS (Park et al., 2017; Song et al., 2015; Wong et al., 2020). We investigated the labeling pattern of a series of 9 to 45 different bifucosylated N-glycans (the number varied depending on cell line). Since each of these N-glycans must have two fucose residues it can either incorporate two residues of ^12^C-fucose (M+0), two residues of ^13^C-UL-fucose (M+12) or one of each (M+6). We expected an increase of label incorporation that is proportional to fucose^Ex^ concentration.

We labeled seven different cell lines with increasing concentrations of exogenous ^13^C-UL-Fucose for 24 h and analyzed the N-glycans produced by these cells. As expected, we observed a dose dependent incorporation of ^13^C-UL-Fucose into N-glycans with a single fucose residue, until we reach a plateau and maximized incorporation of fucose^Ex^ by 24 h (**Figure 3A, Supplementary Figure 4D**). Unexpectedly, for those N-glycans with two fucoses, the first fucose, which we believe is the predominantly the core fucose, requires less fucose^Ex^ to become labeled than the second one. At low fucose^Ex^ concentration (0-10 μM), the first fucose residue originates from fucose^Ex^, while the second uses fucose^Endo^. Replacing both ^12^C-fucoses with ^13^C-UL-Fucose requires more accessible fucose^Ex^ (**Figure 3A, Supplementary Figure 4D**). This suggests fucose in different linkages and added by various fucosyltransferases, localized in different Golgi compartments, relies on distinct GDP-fucose pools. These results are incompatible with a single, homogenous GDP-fucose pool in the cytoplasm, further confirming our hypothesis.

**Figure 3.**
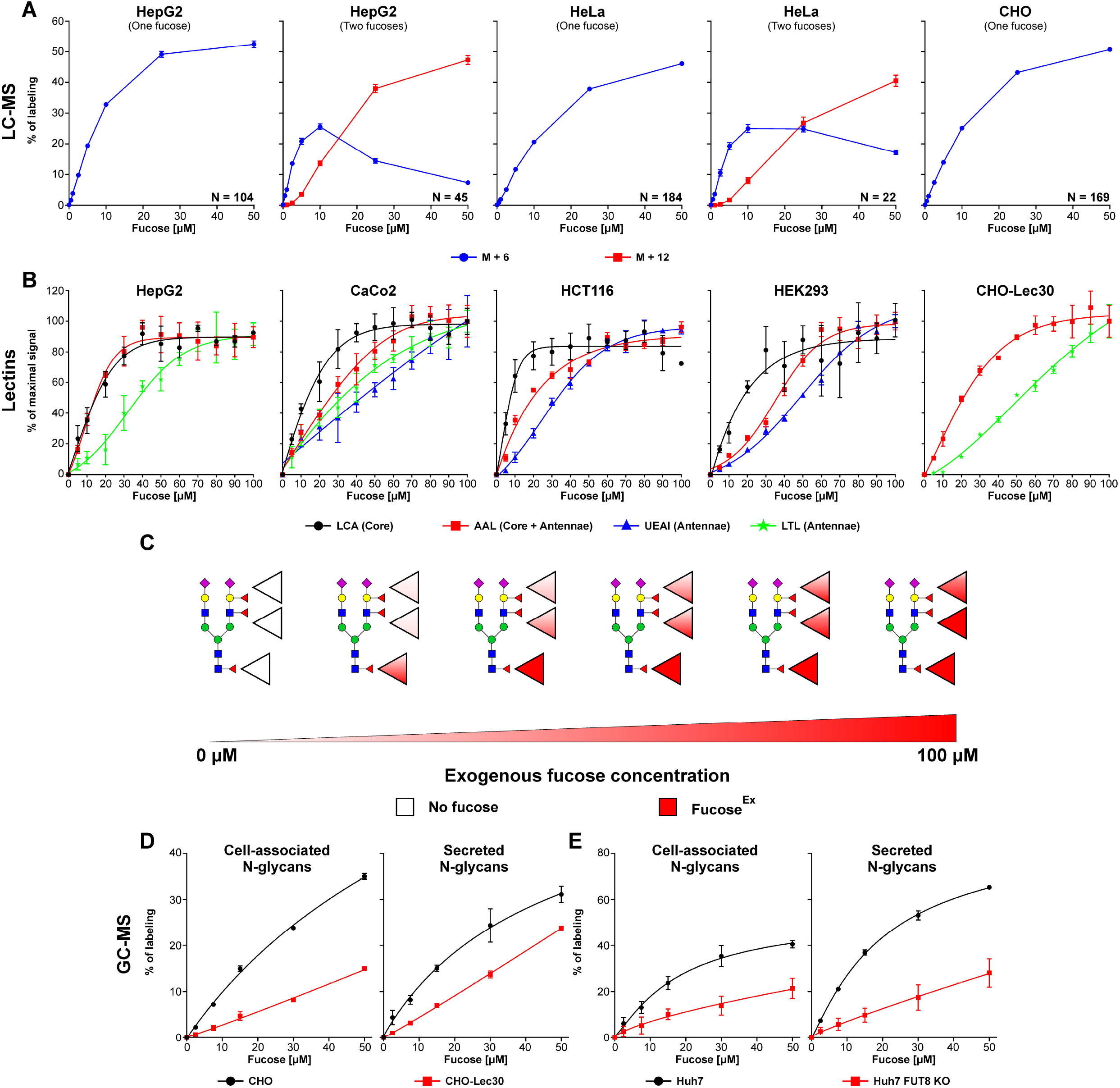
Various fucose linkages exhibit different preference to exogenous fucose. (A) LC-MS analysis of exogenous ^13^C-UL-L-fucose incorporation into cell associated N-glycans produced by HepG2, HeLa and CHO cells that have only one fucose residue as well as N-glycans with two fucoses; M+6 refers to N-glycans which incorporated a single molecule of exogenous fucose; M+12 refers to N-glycans which incorporated two molecules of exogenous fucose; data are presented as mean ± SEM; N refers to the number of unique N-glycan structures. (B) Lectin staining of HepG2, CaCo2, HCT116, HEK293 and CHO-Lec30 cells that have chemically or genetically inactivated *de novo* pathway, treated with increasing concentrations of exogenous fucose; n = 3; data are presented as mean ± SD. (C) Schematic representation of the results presented in panel B. (D) GC-MS analysis of exogenous ^13^C-UL-fucose incorporation efficiency into fucose associated with cellular N-glycoproteins as well as newly synthesized N-glycoproteins secreted by CHO and CHO-Lec30 cells expressed as a percentage of labeling; n = 3; data are presented as mean ± SD. (E) GC-MS analysis of exogenous ^13^C-UL-fucose incorporation into fucose associated with cell associated N-glycoproteins as well as newly synthesized N-glycoproteins secreted by Huh7 and Huh7 FUT8 knock-out cells; n = 3; data are presented as mean ± SD.

Next, we used lectins that specifically recognize fucose in different linkages: LCA recognizes α1-6 fucose (core), LTL recognizes α1-3 and α1-4 fucose, UEAI recognizes α1-2 fucose, and AAL recognizes all these linkages with variable efficiencies. We analyzed incorporation of fucose^Ex^ into different positions in glycans using cells with chemically or genetically inactivated *de novo* pathway (see methods). We chose the cell lines HepG2, CaCo2, HCT116, HEK293 and the CHO mutant Lec30, as they exhibited high binding to LTL and/or UEAI. Once again, we found α1-6 fucose (chitobiose core) requires less fucose^Ex^ to become maximally labeled than α1-2, α1-3 and α1-4 fucose (antennae) (**Figure 3B, 3C**), confirming the results of the above ^13^C-UL-Fucose labeling experiments.

A limitation of the lectin-based approach is that it requires inhibition of the *de novo* pathway and only focuses on fucose^Ex^ utilization. Therefore, we searched for alternative methods to assess incorporation of fucose^Ex^ into different positions of N-glycans, using cells with a functional *de novo* pathway. First, we used GC-MS to compare the incorporation of ^13^C-UL-fucose^Ex^ into N-glycans produced by CHO and CHO-Lec30 cells as well as Huh7 and Huh7 FUT8 knock-out cells. FUT8 is the only fucosyltransferase capable of adding fucose to the chitobiose core of N-glycans (Yang and Wang, 2016) and is the only fucosyltransferase adding fucose to N-glycans, which is expressed in wild-type CHO cells. CHO-Lec30 is a gain of function mutant cell line which additionally expresses FUT4 and FUT9 that fucosylate N-glycan antennae (North et al., 2010; Patnaik and Stanley, 2006). If all fucosyltransferases had similar access to different GDP-fucose sources CHO-Lec30 and Huh7 FUT8 knock-out cells would incorporate comparable amounts of fucose^Ex^ to wild-type CHO and Huh7 cells respectively. However, we found that both CHO-Lec30 and Huh7 FUT8 knock-out cells rely on fucose^Endo^ much more than wild-type CHO and Huh7 respectively, which incorporated significantly higher amounts of fucose^Ex^ to their N-glycans (**Figure 3D, 3E**). This supports our hypothesis on distinct preferences between different GDP-fucose sources among various fucosyltransferases. It also shows that efficient incorporation of fucose^Ex^ into the N-glycan chitobiose core is not due to FUT8 having a better access to GDP-fucose than other fucosyltransferases that modify the N-glycan antennae.

To confirm the linkage specificity, we used an LC-MS approach to analyze N-glycans released from HepG2 secreted proteins labeled with exogenous ^13^C-UL-Fucose (**Supplementary Figure 5)**. Among all applied approaches this is the only method that allowed us to directly investigate fucose linkage. Analysis of three most abundant secreted N-glycans with a single α1-6 fucose, showed a similar trend in fucose^Ex^ incorporation compared to cell associated material analyzed with NanoHPLC Chip-Q-TOF MS (**Figure 4A, Figure 3A**). The same was true when we assessed incorporation of fucose^Ex^ into the two most abundant N-glycans with two fucoses, where the first fucose requires less fucose^Ex^ to become labeled compared to the second one (**Figure 4A**). To investigate which fucose residue is preferentially labeled with fucose^Ex^, we performed MS/MS analysis of bifucosylated N-glycans which only incorporated a single molecule of fucose^Ex^ (**Supplementary Figure 5)**. This allowed us to compare the positions where ^13^C-fucose^Ex^ and ^12^C-fucose^Endo^ are incorporated in N-glycans, determining if there is any preference towards incorporation of either of them into N-glycan chitobiose core vs. antennae. If there was an equal preference of incorporating fucose^Ex^ to N-glycan core and antennae there would be a similar distribution of ^13^C-labled between both fragments of N-glycan. However, we found that fucose^Ex^ is more efficiently incorporated into the chitobiose core, while fucose^Endo^ is preferentially incorporated to the antennae (**Figure 4B, 4C**). Altogether, this again confirms preferential utilization of GDP-fucose originating from the fucose^Ex^ by FUT8, while fucose^Endo^ is used by fucosyltransferases located in the trans-Golgi to modify N-glycan antennae.

**Figure 4.**
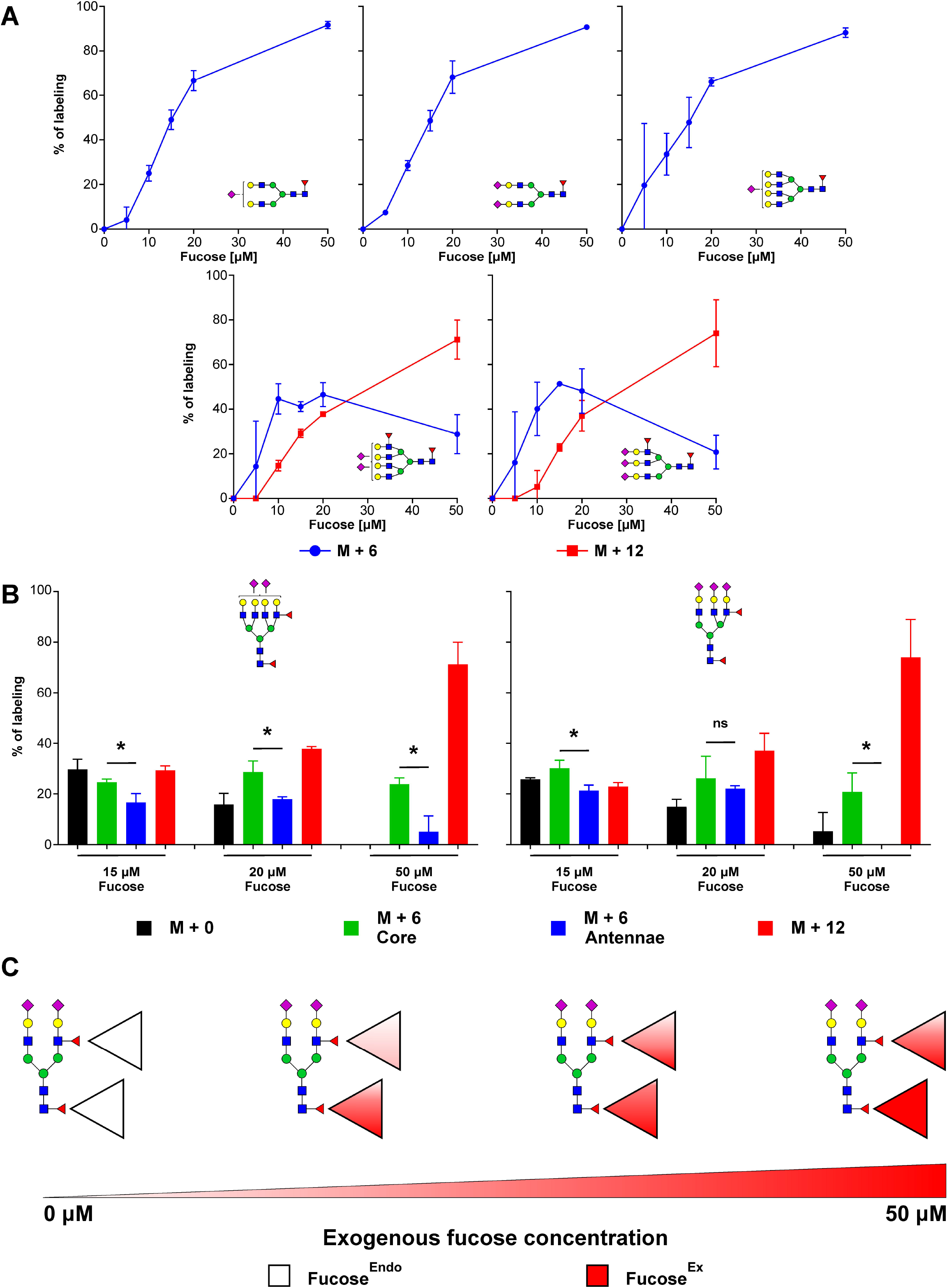
Core fucose preferentially rely on exogenous fucose while fucose attached to N-glycan antennae utilizes endogenous fucose more efficiently. (A) MALDI-TOF analysis of exogenous ^13^C-UL-fucose incorporation into N-glycans that have only one fucose residue as well as N-glycans with two fucose residues, produced by HepG2 cells; M+6 refers to N-glycans which incorporated a single molecule of exogenous fucose; M+12 refers to N-glycans which incorporated two molecules of exogenous fucose; n = 2; data are presented as mean ± SD. (B) LC-MS analysis of exogenous ^13^C-UL-fucose and endogenous ^12^C-fucose incorporation into N-glycans produced by HepG2 cells with two fucose residues. Black bars indicate N-glycans with endogenous ^12^C-fucose incorporated into both chitobiose core and antennae. Green bars indicate N-glycans with exogenous ^13^C-UL-fucose incorporated into chitobiose core and endogenous ^12^C-fucose into antennae. Blue bars indicate N-glycans with exogenous ^13^C-UL-fucose incorporated into antennae and endogenous ^12^C-fucose into chitobiose core. Red bars for N-glycans with exogenous ^13^C-UL-fucose into both chitobiose core and antennae; n = 2; data are presented as mean ± SD; statistical analysis was performed using *t*-test; p > 0.1 ns; p < 0.1 *. (C) Schematic representation of the results presented in panel B.

### Different glycoprotein acceptors rely on distinct GDP-fucose sources

Next, we characterized the contribution of different GDP-fucose sources into O-fucosylation of Epidermal Growth Factor (EGF)-like repeats and ThromboSpondin type 1 Repeats (TSRs), which occur in the ER and are performed by GDP-fucose protein O-fucosyltransferases (POFUTs). The former reaction is catalyzed by POFUT1, while the latter by POFUT2 (**Figure 5A**). Both enzymes catalyze chemically identical reactions and only differ in recognized acceptors/glycoproteins. Over 100 proteins, including Notch, contain the EGF-like repeat consensus sequence required for POFUT1 O-fucosylation and approximately 50 proteins contain a TSRs consensus sequence for POFUT2 O-fucosylation (Schneider et al., 2017).

**Figure 5.**
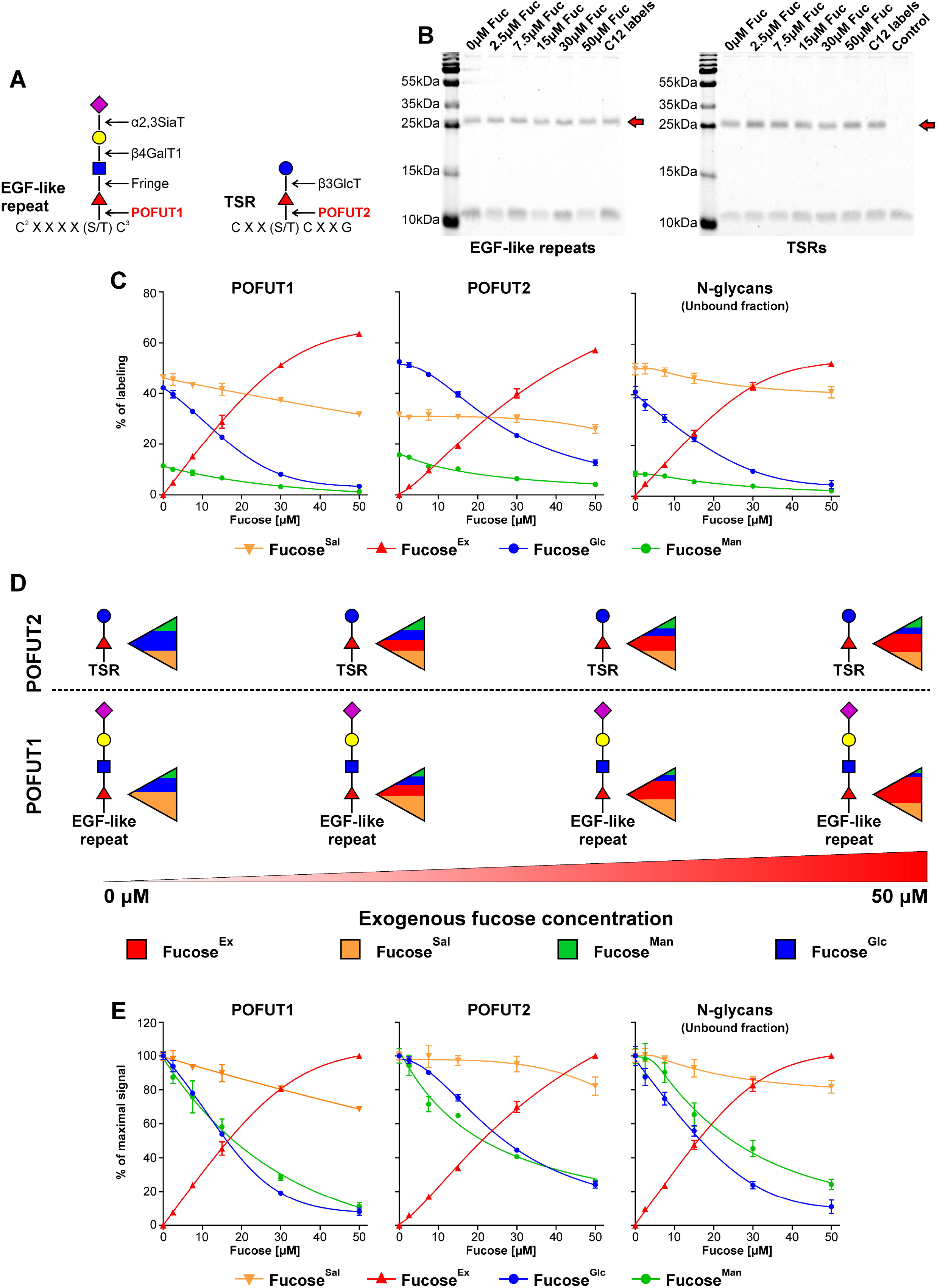
Various fucosyltransferases, which fucosylate different glycoproteins, exhibit distinct preference to different GDP-fucose sources. (A) Schematic showing POFUT1 and POFUT2 dependent glycosylation of EGF-like repeats and TSRs. (B) Coomassie Brilliant Blue G250 staining of a representative SDS-PAGE gel of purified recombinant EGF-like repeats and TSRs secreted for 12 h by Huh7 cells. Red arrows indicate the protein of interest. (C and E). Incorporation of 5 mM ^13^C-1,2-glucose, 50 μM ^13^C-1,2,3-mannose and ^13^C-6-fucose into fucose associated with EGF-like repeats or TSRs secreted by Huh7 cells pre-labeled for eleven days with 50 μM ^13^C-UL-fucose expressed as a percentage of labeling (C) and a percentage of maximal signal (E); n = 2 (for POFUT1 and POFUT2) and n = 4 (for unbound N-glycans); data are presented as mean ± SD. (D) Schematic representation of the results presented in panel C.

We used Huh7 cells pre-labeled with ^13^C-6-fucose overexpressing recombinant proteins containing either NOTCH1 EGF-like repeats 1-5, where repeats 2, 3 and 5 have a single O-fucose modification, or a recombinant protein containing thrombospondin 1 TSRs 1-3, where all three TSRs carry an O-fucose modification. These proteins were purified using affinity cobalt resin and their purity was confirmed in SDS-PAGE (**Figure 5B**). If both POFUTs had the same access to various GDP-fucose pools they would rely on them to a comparable extent. However, we found that POFUT1 relies more on GDP-fucose originating from a fucose^Ex^ as well as fucose^Sal^, compared to POFUT2, which has a stronger preference for GDP-fucose from the *de novo* pathway (**Figure 5C, 5D**). In addition, we did not observe any difference in a sensitivity of fucose^Man^ and fucose^Glc^ to fucose^Ex^ as was the case of fucosylated N-glycoproteins. For POFUT1 and POFUT2, these sources are treated comparably (**Figure 5E**). When analyzing secreted material that did not bind to the affinity cobalt resin, we found that overexpression of EGF-like repeats or TSRs does not alter normal fucosylation of N-glycans (**Figure 5C, 5E**). This experiment also demonstrated that N-glycan-linked fucose relies on various GDP-fucose pools to a different extent compared to O-linked fucose. Altogether, the data shows that different fucosyltransferases and various glycoproteins utilize distinct GDP-fucose sources, and confirms our hypothesis on the presence of separate, non-homogenous GDP-fucose pools in the cytoplasm.

## DISCUSSION

The results presented here support our hypothesis that the metabolic origin of fucose determines its use for glycosylation and suggests that GDP-fucose resides in multiple, distinct pools. We can simultaneously distinguish the contributions of exogenous, salvaged and *de novo* sources to global N-glycosylation and that of individual glycoproteins. It is broadly accepted that these sources contribute to a single, homogenous pool and little is known about the interplay between these pathways. There are multiple reports on biosynthesis of nucleotides and lipids that compare the contribution of the *de novo* biosynthesis to exogenous precursors, but they never simultaneously address all biosynthetic routes (Moitra et al., 2021; Nieto et al., 2008; Wu et al., 2020; Zhang et al., 2019).

The pioneering studies performed in HeLa cells that concluded 90 % of GDP-fucose is derived from the *de novo* pathway used only a single fucose^Ex^ concentration, 0.3 μM (Yurchenco and Atkinson, 1975, 1977). In a dose-response experiments, we showed fucose^Ex^ is the preferential source of N-glycan-associated fucose, gradually suppressing the *de novo* pathway without increasing the total cytoplasmic concentration of GDP-fucose. It agrees with previous experiments in HepB3 cells showing that 100 μM fucose^Ex^ does not increase GDP-fucose (Moriwaki et al., 2007). Suppression of the *de novo* pathway is likely due to feedback inhibition of GMDS with GDP-fucose, previously showed for human and *E.coli* enzyme (Kornfeld and Ginsburg, 1966; Sullivan et al., 1998). We also looked at the contribution of fucose^Man^ and fucose^Glc^ into N-glycan-associated fucose. Since increasing concentrations of fucose^Ex^ do not affect glucose and mannose contribution into N-glycan-associated mannose, we would expect that fucose^Ex^ affects them equally. However, that is not the case and we observe that fucose^Man^ is less sensitive to fucose^Ex^ than fucose^Glc^. It indicates that different, non-homogenous pools of both GDP-fucose and GDP-mannose may exist in the cytoplasm.

Little is known about monosaccharides salvage since most studies equate it with exogenous substrate. Analysis of N-acetylneuraminic acid (Neu5Ac) recycling using cells from individuals with Salla disease (Aula et al., 2000), which characterize with dysfunctional SLC17A5 lysosomal transporter and are unable to recycle Neu5Ac (Ghosaini et al., 1988), demonstrated the importance of this process in sialylation (Chigorno et al., 1996). A single study using a lysosomal inhibitor NH_4_Cl acidifying lysosomal lumen and causing a disfunction of glycosydases, demonstrated that N-acetylglucosamine (GlcNAc) and N-acetylgalactosamine (GalNAc) are also efficiently reutilized from glycoconjugates (Rome and Hill, 1986). We investigated the contribution of fucose^Sal^ into fucosylation, while simultaneously analyzing its interplay with other GDP-fucose sources. In absence of fucose^Ex^ ~50 % of N-glycan-associated fucose originated from fucose^Sal^ and another ~50 % came from the *de novo* pathway. In contrast to the *de novo* pathway, salvage is almost completely insensitive to fucose^Ex^, and its contribution is quite stable in its increasing concentrations. Since the total cytoplasmic concentration of GDP-fucose does not change much upon the exposition to fucose^Ex^ there has to be a mechanism regulating a contribution of fucose^Sal^. However, further studies are needed to understand how fucose^Ex^ selectively inhibits the salvage pathway as both sources of GDP-fucose utilize the same biosynthetic machinery.

Fucosylation of N-glycan chitobiose core is catalyzed by FUT8 and occurs in the early medial-Golgi (Stanley, 2011), while antenna fucosylation happens in the trans-Golgi as well as trans-Golgi network (Brito et al., 2008; Buffone et al., 2013; Sousa et al., 2004). Fucose^Ex^ is more efficiently incorporated to the N-glycans chitobiose core than into antenna. Since the addition of fucose^Ex^ does not change the size of GDP-fucose pools, this observation cannot be explained by differences in a K_m_ between distinct fucosyltransferases. Analysis of FUT8 knock-out cells disproves that the preference of fucose^Ex^ by FUT8 is simply due to its utilization of the GDP-fucose pool before it becomes accessible to other fucosyltransferases (i.e FUT4, FUT9). If that were true, elimination of FUT8 would make the GDP-fucose originating from fucose^Ex^ more accessible to other fucosyltransferases. In contrast, it decreases the overall utilization of fucose^Ex^. We believe that separate Golgi compartments have different access to various GDP-fucose pools.

In the ER POFUT1 and POFUT2 add a single O-fucose residue to EGF-like repeats and TSRs respectively (Schneider et al., 2017). O-fucosylation of a disintegrin and metalloproteinase with thrombospondin motifs (ADAMTS) proteins performed by POFUT2 is required for their secretion as it was shown for ADAMTS9, ADAMTS13 and ADAMTS20 (Benz et al., 2016; Holdener et al., 2019; Ricketts et al., 2007). O-fucosylation of Notch, conducted by POFUT1, is required for activation of Notch by its ligand (Shi and Stanley, 2003). This shows the importance of O-fucosylation in a variety of biological processes. We looked at the utilization of various GDP-fucose pools by POFUT1 and POFUT2, demonstrating that POFUT1 exhibits significantly higher preference towards exogenous and salvaged substrate, POFUT2 more efficiently utilizes *de novo* produced GDP-fucose. It indicates that fucosyltransferases for O-glycans located outside the Golgi and modifying distinct glycoproteins also rely on separate GDP-fucose pools. The only possible explanation for this observation is an existence of separate, non-homogenous GDP-fucose pools in the cytoplasm. It is highly probable that the ER and Golgi have a differential access to the GDP-fucose originating from various sources. This would also be true in case of distinct Golgi stacks as we observe that FUT8, located in the early medial Golgi has a stronger preference towards fucose^Ex^ comparing to the fucosyltransferases occupying trans Golgi stacks. Altogether, it suggests that different GDP-fucose pools are accessible to the ER, medial Golgi and trans Golgi. How separate pools would be generated and maintained in the cells remains unclear. We do not believe that they are separated from each other through their allocation inside a membrane-delimited structure. Our hypothesis is that monosaccharides have distinct ways of cell entry and handling within the cell, contributing to the GDP-fucose pool segregation. Furthermore, each pathway utilizes various enzymes for GDP-fucose biosynthesis that could support nucleotide sugar segregation. In this scenario, each pool exists due to an increase in local GDP-fucose concentration and is variably accessible to a limited number of fucosyltransferases and protein acceptors. We suggest that GDP-fucose biosynthetic enzymes form functional complexes and their association as well as their subcellular distribution is influenced by fucose^Ex^ concentration. However, it is unclear how it could regulate this process.

Study on exogenous N-azidoacetylglucosamine (GlcNAz) incorporation into N-glycans showed that this GlcNAc analogue is efficiently added into the antennae, but not into chitobiose core (Shajahan et al., 2020). The former process occurs in the Golgi and the latter on the ER membrane. It suggests that there is a preference of one source of UDP-GlcNAc over the other during different stages of N-glycan biosynthesis. Recent studies on glycogen metabolism in brain demonstrated that it serves as a reservoir for multiple glycoconjugates. Disruption of its metabolism causes global decrease in free pools of UDP-GlcNAc and N-glycosylation (Sun et al., 2021). These studies are consistent with our proposed model of distinct pools of the same nucleotide sugar existing in the cytosol, where one pool is preferred over another by different glycosyltransferases.

To summarize, we presented a novel concept of selective utilization of different monosaccharide sources using fucose as a model sugar. Our research demonstrates that there are separate, non-homogenous pools of nucleotide sugars in the cytoplasm, and each makes a specific contribution to glycosylation processes. Various glycoproteins and distinct glycosyltransferases rely on different nucleotide sugar pools and this selectivity is regulated by the access of exogenous/dietary monosaccharides. We think that it is worthwhile to expand this concept beyond glycosylation and verify it for other metabolic precursors.

## ACKNOWLEDGMENT

The Rocket Fund and National Institutes of Health (NIH) grant R01DK099551 awarded to H.H.F., GM061126 awarded to R.S.H., S10OD018530 and R24GM137782 awarded to P.A. and R01GM049077 awarded to C.B.L. supported this work. P.S. was supported by Frontiers in Congenital Disorders of Glycosylation Career Developmental Award U54 NS115198. The Sanford Burnham Prebys Cancer Metabolism Core was supported by Cancer Center Support grant P30CA030199 from the National Cancer Institute. We would like to thank Dr. Pamela Stanley for providing with CHO, CHO-Lec13 and CHO-Lec30 cells. We would like to thank Jamie Smolin for her assistance. We would like to thank Amgen for providing Fucostatin II. We would like to thank Dr. Jeffrey Esko, Dr. Robert Coffman, Dr. Paul Atkinson, and Dr. Christopher Newgard for their critical reviews.

## AUTHOR CONTRIBUTIONS

Conceptualization, P.S., and H.H.F.; methodology, P.S., B.G.N., L.E.P., A.S., M.W., D.A.S., K.M., Z-J.X., C.B.L., R.S.H., P.A., and H.H.F.; investigation, P.S., B.G.N., L.E.P., A.S., M.W., D.A.S., K.M., and Z-J.X.; supervision, C.B.L., R.S.H., P.A., and H.H.F.; writing – original draft, P.S., H.H.F., and B.G.N.; writing – review & editing, all authors.

## MATERIALS AND METHODS

### Cell culture

HepG2, Huh7, HEK293, HeLa, HCT116, CaCo2 and A549 (ATCC) cells were grown in complete 1 g/l (5.5 mM) glucose containing Dulbecco’s modified Eagle’s medium (Corning) supplemented with 10 % heat-inactivated fetal bovine serum (Sigma-Aldrich), 100 U/ml penicillin and 10 mg/ml streptomycin (GIBCO) and 2 mM L-glutamine. CHO, CHO-Lec13 and CHO-Lec30 were grown in complete Ham’s F-12K (Kaighan’s) medium with 7 mM glucose (Corning) supplemented with 10 % heat-inactivated fetal bovine serum, 100 U/ml penicillin and 10 mg/ml streptomycin and 2 mM L-glutamine. For metabolic labeling purposes cells were grown in a glucose free Dulbecco’s modified Eagle’s medium (Gibco) supplemented with 5 mM glucose, 50 μM mannose and 0-50 μM fucose, 10 % heat-inactivated fetal bovine serum, 100 U/ml penicillin and 10 mg/ml streptomycin and 2 mM L-glutamine. Glucose, mannose and fucose were variously replaced with equimolar ^13^C substrates as noted in the next section. In the case of CHO, CHO-Lec13 and CHO-Lec30 labeling medium was additionally supplemented with 0.6 mM L-proline as these cell lines are auxotrophic for proline. In studies on secreted N-glycans or glycoproteins cells were grown in serum free medium.

### Metabolic labeling

To analyze the contribution of different monosaccharides into glycosylation, cells were grown in different combinations of ^13^C-glucose, ^13^C-mannose and ^13^C-fucose. In each experiment, the number of heavy carbons varied between glucose, mannose and fucose substrates. The following sugars were used in this study: ^13^C-5-glucose (M+1), ^13^C-1,2-glucose (M+2), ^13^C-UL-glucose (M+6), ^13^C-4-mannose (M+1), ^13^C-3,4-mannose (M+2), ^13^C-1,2,3-mannose (M+3), ^13^C-UL-mannose (M+6), ^13^C-6-fucose (M+1) and ^13^C-UL-fucose (M+6). To analyze fucose salvage, cells were pre-labeled with either ^13^C-6-fucose (M+1) or ^13^C-UL-fucose (M+6) by growing them for eleven days in complete medium containing 50 μM ^13^C-fucose. This allowed for substitution of 90-95 % of N-glycan-associated fucose with ^13^C-fucose. To distinguish between salvaged and exogenous fucose, fucose^Sal^ carried a different number of heavy carbons comparing to fucose^Ex^. All ^13^C-labeled monosaccharides were purchased from Omicron Biochemicals.

### GC-MS analysis and sample preparation

Cells were labeled with ^13^C-sugars for 12 h, washed twice with DPBS, scraped, and centrifuged for 3 min at 1,600 rpm. Cell pellets were sonicated in 50 mM sodium phosphate buffer pH 7.5 containing 40 mM DTT, incubated at 100 °C, cooled down and sonicated again. PNGase F was added to the resulting cell lysate and samples were incubated overnight at 37 °C. For analysis of secreted N-glycans HepG2 cells were labeled for 6 h before collecting the medium, while Huh7 and CHO were labeled for 12 h. Serum free medium was concentrated using 3 kDa cut-off filters (Sigma-Aldrich). Next, buffer was changed to 50 mM sodium phosphate buffer pH 7.5 using the same filters. Concentrated medium was collected from the filters and, after addition of PNGase F, samples were incubated overnight in 37 °C.

Released N-glycans were purified with TopTip 10-200 μl carbon columns (PolyLC Inc.). All columns were prewashed using 1 × 200 μl 80 % acetonitrile solution with 0.1 % TFA, 1 × 200 μl 50 % acetonitrile solution with 0.05 % TFA and 1 × 200 μl 3 % acetonitrile solution with 0.05 % TFA, followed by equilibration using 2 × 200 μl water. Each time the solution was removed from the columns by centrifuging for 5 min at 5,000 rpm. Cell samples were centrifuged for 20 min at 14,000 rpm and the resulting supernatant was loaded onto the columns. Medium samples were directly loaded onto the columns. After loading the samples, columns were subsequently washed twice with 200 μl of water and twice with 200 μl of 3 % acetonitrile solution with 0.05 % TFA. Glycans were eluted with 3 × 200 μl 50 % acetonitrile solution with 0.05 % TFA and dried in a SpeedVac.

Dried N-glycans were hydrolyzed in 2 N TFA (Sigma Aldrich) at 100 °C for 2 h to release monosaccharides and cooled samples were again dried in a SpeedVac. If both fucose and mannose were analyzed, the hydrolyzed sample was equally divided. Samples were stored in −20 °C and derivatized immediately before GC-MS analysis.

To analyze the contribution of glucose and mannose into N-glycan-associated mannose, hydrolyzed samples were dissolved in 10 μl water and 50 μl of 50 mg/ml hydroxylamine hydrochloride (Sigma Aldrich) in 1-methyl-imidazole (Sigma Aldrich) was added. Samples were incubated at 80 °C for 10 min. After cooling for 10 min, 100 μl of acetic anhydride (Sigma Aldrich) was added to the samples, followed by 100 μl of chloroform (Honeywell) and subsequent addition of 200 μl of water. Samples were vigorously vortexed for 10-15 s and centrifuged for 10 min at 14,000 rpm. Next, the top layer (aqueous) was discarded and 200 μl of water was added to the organic phase. Samples were again mixed, centrifuged and the aqueous layer discarded. The organic layer was dried in a SpeedVac, resuspended in pyridine (Chem-Impex int’l Inc.), and analyzed with GC-MS. GC-MS analysis was done using a 15 m × 0.25 mm × 0.25-μm Rxi-5ms column (Restek) on a QP2010 Plus GC-MS. Detector sensitivity and m/z rage were optimized for each sample set separately. Electron impact was used for MS ion fragmentation. GC-MS ion fragment intensities were quantified using GC-MS Solution version 2.5o SU3 from Shimadzu Corp. Each fragment was corrected for the natural abundance of each element using matrix-based probabilistic methods as described (Nanchen et al., 2007; van Winden et al., 2002). Contribution of glucose and mannose into mannose was calculated from the fragment m/z 314 (Ichikawa et al., 2014).

To analyze the contribution of glucose, mannose and fucose into N-glycan-associated fucose hydrolyzed samples were derivatized initially in 20 mg/ml pyridine solution of O-(2,3,4,5,6-Pentafluorobenzyl) hydroxylamine hydrochloride (PFBO; Alfa Aesar) and incubated for 20 min at 80 °C. After cooling, an equal volume of N,O-Bis(trimethylsilyl)trifluoroacetamide (BSTFA; Sigma Aldrich) was added and samples were incubated for additional 60 min at 80 °C. Derivatized samples were analyzed by GC-MS using the instrument and column described above, but with negative chemical ionization, using methane as the reagent gas. The GC-MS was programmed with an injection temperature of 250°C, 2 μl injection volume and split ratio 1/10. The GC oven temperature was initially 160°C for 4 min, rising to 230°C at 6°C/min, then to 280°C at 60°C/min with a final hold at this temperature for 2 min. GC flow rate with helium carrier gas was 50 cm/s. The GC-MS interface temperature was 300°C and (NCI) ion source temperature was 200°C. Detector sensitivity and m/z range were optimized for each sample set separately. Contribution of glucose, mannose and fucose into fucose was calculated from the fragments with m/z 376 and 466 (respectively, C_15_H_34_O_4_NSi3 – loss of pentafluorobenzene plus trimethylsilyl-OH, and C_18_H_44_O_5_NSi_4_ – loss of pentafluorobenzene) (**Supplementary Figure 1A, 1B, 1C**). Same as for mannose, GC-MS ion fragment intensities were quantified using GC-MS Solution version 2.5o SU3 from Shimadzu Corp and each fragment was corrected for the natural abundance of each element using matrix-based probabilistic methods as described (Nanchen et al., 2007; van Winden et al., 2002). To ensure the accuracy of the data analysis process we always performed control experiments with cells growing in 5 mM ^12^C-glucose and 50 μM ^12^C-mannose and compared the isotopic distribution on M+0 to M+6 positions between ^12^C- and ^13^C-labled samples. In addition, we used HCT116 and CHO-Lec13 cells, which are GMDS null mutants that do not produce GDP-fucose *de novo,* therefore do not utilize glucose or mannose for fucosylation. We did not see a significant contribution of glucose metabolism products into fucosylation.

### S^35^ methionine/cysteine labeling

HepG2 cells were seeded on 6 cm plates and at 70 % confluency were washed twice with DPBS. Next, labeling complete Dulbecco’s modified Eagle’s medium without methionine (Corning) supplemented with 0.2 mCi S^35^ Met/ml (1000 Ci/mmol; 3:1 mix of Met and Cys; American Radiolabeled Chemicals, Inc.; cat #ARS 0110A) was added. After 2 h of labeling, cells were washed 5 times with DPBS, and medium was replaced with serum free medium containing 10 fold excess of cold Met (300 mg/l). Medium was collected after 0 min, 15 min, 30 min, 1 h, 1.5 h, 2 h, 3 h or 6 h of incubation. To analyze label incorporation into proteins, 100 μl of medium was mixed with 200 μg of bovine serum albumin. An equal volume of 20 % TCA was added to each sample and incubated on ice for 20 min then centrifuged at 14,000 rpm for 15 min at 4 °C. Precipitated proteins were washed twice with 1 ml of ice-cold acetone and centrifuged at 14,000 rpm for 15 min at 4 °C after each wash. Excess acetone was evaporated by incubating the samples for 1 h in room temperature under ambient atmosphere. To reconstitute proteins, 100 μl of 2 % SDS was added, and samples were sonicated and incubated for 15 min in 100 °C. 10% of resulting solution was counted in a scintillation counter.

### [5, 6-^3^H]-fucose labeling

HepG2 cells were seeded on 6-well plates and at 70 % confluency, they were washed twice with DPBS and labeling complete Dulbecco’s modified Eagle’s medium containing 5 μM cold fucose and 15 μCi of L-[5,6-^3^H]-fucose (40-60 Ci/mmol; 1 mCi/ml; American Radiolabeled Chemicals; cat #ART0106A) per 0.5 ml of the medium was added. After 2 h, cells were washed 5 times with DPBS, and medium was replaced with serum free medium containing 5 μM cold fucose. Medium and cells were collected after 0 min, 15 min, 30 min, 60 min, 120 min or 180 min of incubation. Before collecting cells, they were washed 5 times with DPBS, scraped and lysed in 2 % SDS. 50% of collected material was counted using a scintillation counter.

### LCA pull down

LCA agarose beads (Vector Laboratories) were equilibrated by washing 5 times with 10 mM HEPES buffer pH 7.5 with 0.15 M NaCl 0.1 mM CaCl2 and 1 mM MnCl2. Next, 100 μl of medium sample was diluted with 900 μl of equilibration buffer and incubated with 30 μl of a beads overnight with rotation at 4 °C. Subsequently, the beads were washed 5 times with equilibration buffer. Between each step, samples were centrifuged for 5 min, 1,000 × g, 4 °C and the fractions were saved for further analysis. To elute fucosylated N-glycans, 200 μl of 2 % SDS was added to the beads and samples were heated at 100 °C for 10 min. The solution was then centrifuged for 15 min at 14,000 rpm and transferred to a fresh tube. 10 % of each fraction was counted in a scintillation counter.

### FUT8 knock-out in Huh7 cells

The guide RNA sequence CAGAATTGGCGCTATGCTAC targeting exon 7 of *FUT8* was cloned into px458 pSpCas9(BB)-2A-GFP (Addgene). The resulting plasmid was transfected into Huh7 cells using ViaFect (Promega) transfection reagent following the manufacturer’s instructions. 48 h post-transfection, GFP positive cells were FACS sorted using a FACS ARIA IIu instrument (BD Biosciences) into 96 well plates to generate isogenic clones, which were expanded and screened by Sanger sequencing with forward primer GCTGGTGTGTAATATCAACA and reverse primer GTATGTTCCATGAAGGTCTAC. Two isogenic clones were identified carrying different homozygous mutations in *FUT8* (NM_178155.2). Clone 3 harbored a c.732_762del [p.E244Dfs*9] and clone 20 a c.753dupT [p.T252Yfs*9]. FUT8 deficiency was confirmed in Western Blot using a Fut8-specific antibody (Proteintech). The glycosylation defect was confirmed with LCA and AAL lectins (Vector Laboratories). Clone 3 was used in further analysis.

### Isolation and quantitative HPLC analysis of nucleotide sugars

To measure the amount of GDP-fucose in cells, cytoplasmic nucleotide sugars were isolated using a modified protocol described by Nakajima et al. (Nakajima et al., 2010). 500 pmol of UDP-arabinose, an internal standard, was added prior to workup. We chose UDP-arabinose since it does not occur in mammalian cells and has a unique retention time in the elution gradient (**Supplementary Figure 2A**). Linear response of authentic GDP-fucose was confirmed between 40 and 500 pmol (**Supplementary Figure 2B**).

Briefly, cells were grown on 10 cm plates to 70-80 % confluency. For the last 6 h (HepG2) or 12 h (Huh7 and CHO) they were grown in a presence or absence of 50 μM fucose. Cells were scraped in DPBS and centrifuged for 3 min at 1,500 rpm. Resulting cell pellet was washed twice with cold DPBS. Next, 1 ml of 70 % ice-cold ethanol containing 500 pmol of UDP-arabinose was added to the cell pellet. Cells were sonicated and centrifuged at 16,000 × g for 10 min at 4 °C to remove insoluble material. Supernatant was lyophilized and pellet used to determine protein concentration using BCA Protein Assay Kit (Pierce). Dried supernatant was dissolved in 1 ml of 5 mM ammonium bicarbonate and purified on a 3 ml porous graphitic carbon column (EnviCarb; Sigma Aldrich – Supelco) equilibrated with 2 ml of 80 % acetonitrile solution with 0.1 % TFA and 2 ml of water. After loading the sample, the column was sequentially washed: 2 ml of water, 2 ml of 25 % acetonitrile solution, 2 ml of water, 2 ml of 50 mM triethylammonium acetate buffer pH 7.0 and 2 ml of water. Nucleotide sugars were eluted with 2 ml of 25 % acetonitrile solution in 50 mM triethylammonium acetate buffer pH 7.0 and lyophilized. Samples were stored in −80 °C and analyzed within 12-48 h after the isolation.

Nucleotide sugars were separated using Inertsil ODS-4 250 × 4.6 mm column with a particle size 3 μm (GL Science) and Vanquish Flex HPLC system (Thermo Fisher Scientific), with 100 mM potassium phosphate buffer pH 6.4 containing 8 mM tetrabutylammonium hydrogen sulphate as buffer A and 70 % solution of buffer A in 30 % acetonitrile as buffer B. The separation method was as follows: buffer A for 13 min, linear gradient of 100-33 % buffer A for 22 min, linear gradient of 33-0 % buffer A for 1 min, 100 % B for 14 min and linear gradient of 0-100 % A for 1 min. Flow rate was 0.8 ml/min and the separation temperature was 23 °C. Nucleotide sugars were detected using an UV detector and absorption λ_254_.

### NanoHPLC Chip-Q-TOF MS analysis

HepG2, CHO, CHO-Lec30, CaCo2, HeLa, HEK293, A549 and HCT116 cells were labeled for 24 h with increasing concentrations of ^13^C-UL-Fucose, washed 3 times with DPBS, scraped and centrifuged for 3 min at 1,600 rpm. Resulting cell pellets were stored in −80 °C. N-glycans were isolated and analyzed as described by Wong et al. (Wong et al., 2020). Cell pellets were sonicated in 20 mM HEPES buffer pH 7.5 containing 0.25 M sucrose and 1:100 protease inhibitor cocktail set V (Calbiochem) and centrifuged at 2,000 × g for 10 min to pellet nuclear fraction. Next, lysates were centrifuged in an ultracentrifuge at 200,000 × g for 45 min. Resulting pellets were dissolved in 100 μl of 100 mM phosphate buffer pH 7.0 containing 5 mM DTT and incubated in 100 °C for 1 min to denaturate proteins. To release N-glycans 2 μl of PNGase F (New England Biolabs) was added and the samples were incubated for 18 h in 37°C. After that they were ultracentrifuged at 200,000 × g for 45 min and supernatants containing free N-glycans were purified on porous graphitized carbon columns and dried in SpeedVac.

Agilent 6520 Accurate Mass Q-TOF LC/MS system with a microfluid porous graphitized carbon chip (Agilent) was used to analyze N-glycans composition. The binary gradient consisted of 0.1 % of formic acid with 3 % acetonitrile in water as buffer A and 1 % formic acid with 89 % acetonitrile in water was used to separate samples. The separation method was as followed: 100 % of A for 2.5 min, linear gradient of 0-16 % B for 17.5 min and linear gradient of 16-58 % B for 15 min, at the constant flow rate of 0.3 μl/min. For the data acquisition the Q-TOF MS was set to acquire only MS1 in positive ionization mode with a cycle time of 1.5 s. To assist with compound identification, control samples were also run with collision-induced dissociation using nitrogen gas. Four MS2 spectra were obtained for every MS1 through data-dependent acquisition, with a total cycle time of 5.25 s. The mass range was set at 600–2000 m/z. The instrument was calibrated with the ESI tuning mix commercially available from Agilent.

To analyze the data, we first identified peaks with Agilent MassHunter B.07 software using existing library described by Wong et al. (Wong et al., 2020). From the exported data we created a new library containing the chemical formulae and retention times of identified N-glycans for targeted isotope extraction. We used the Batch Isotopologue Analysis function of Agilent Profinder B.08 software to extract quantitative isotopologue data for each fucosylated N-glycan we identified. The program averages the MS signals across a chosen peak in an extracted ion chromatogram to form the MS spectra from which isotopologue signals are extracted. It also corrects for the natural distribution of ^13^C and returns the relative abundance of each isotopologue. Mass tolerance was set at ± (10 ppm + 2 mDa) to screen out background noise or signals from other co-eluting compounds. For each glycan, the percent abundance of each isotopologue signal from M + 0 to M + 6 or M +12 was exported for further analysis.

### High throughput fucosylation analysis with lectins

Prior to the experiment, HepG2, CaCo2, HEK293 and CHO-Lec30 cells were grown for 48 h in the presence of 20 μM Fucostatin II (Amgen), which is a small molecule inhibitor of GMDS (**Figure 1A**) and blocks the *de novo* production of GDP-fucose (Allen et al., 2016). Preincubated cells were seeded on 96-well black flat bottom plates (Falcon) in the presence of 20 μM Fucostatin II. After 32 h medium was replaced with fresh medium containing increasing concentrations of fucose and 20 μM Fucostatin II. Cells were incubated with fucose for another 16 h and analyzed by immunofluorescence staining as described in the next paragraph. The experimental procedure was identical for HCT116 cells, except Fucostatin II was not added since these cells are GMDS-deficient and lack a functional *de novo* pathway.

Cells were washed twice with DPBS, incubated for 15 min with 4 % paraformaldehyde, washed three times with DPBS and blocked with Carbo Free Blocking Solution (Vector Laboratories) in DPBS. After 1 h of blocking, biotinylated lectins (Vector Laboratories) were diluted in blocking solution containing 1 mM MgCl_2_, 1 mM CaCl_2_ and 1 mM MnCl_2_ and added for 1 h. LCA and AAL were diluted 1:500 while LTL and UEAI were diluted 1:100. Cells were washed three times with DPBS and incubated for 1 h with Cy3-labeled streptavidin (Vector Laboratories) diluted 1:100 (LTL and UEAI) or 1:500 (LCA and AAL) in blocking solution. DAPI (Thermo Fisher Scientific) was then added directly to the solution, at a final dilution 1:1000. After 20 min, cells were washed three times in DPBS and 100 μl of DPBS was added to each well and plates were sealed with adhesive plate tape (Thermo Fisher Scientific).

Cells were analyzed immediately using IC200 microscope (Vala Sciences) and an air 20× objective. For each well data was collected from 4-6 different focal areas. For data analysis Acapella software (PerkinElmer) was used. The data were expressed as an average ratio of Cy3 fluorescence intensity to DAPI fluorescence intensity for at least 500 cells/well.

### MALDI-TOF-MS and LC-MS/MS analysis

HepG2 cells were washed 6 times with DPBS and grown for 6 h in serum free medium with increasing concentrations of UL-^13^C-fucose. Medium was collected and concentrated using 3 kDa cut-off filters and repeatedly exchanged with 50 mM sodium phosphate buffer pH 7.5 using the same filters. Concentrated medium was collected from the filters and 3 μl of PNGase F (New England Biolabs) was added to each sample then incubated at 37 °C for 24 h. Following incubation, samples were cooled to room temperature and passed through a 10 kDa Amicon Ultra centrifuge filters at 16,000 × g for 15 min. 450 μl of 50 mM ammonium bicarbonate was added to the filters, and samples were again centrifuged. Flow-through was collected and passed through a C18 SPE cartridge (Resprep). N-glycans were eluted with 3ml of 5 % acetic acid and lyophilized.

Per-O-methylation was performed using a dimethylsulfoxide (DMSO)/ sodium hydroxide (NaOH) base. Base was made by combining 100 μl 50 % NaOH and 200 μl methanol in a glass tube. 4 ml of anhydrous DMSO was added to the solution and mixed vigorously. The solution was then centrifuged at 3,000 × g for 5 min. White precipitated solid formed on the top of the solution as well as all remaining DMSO was removed. This process was repeated until the white precipitate no longer formed. The resulting base gel material at the bottom of the tube was then mixed with 1 ml of DMSO. Samples were reconstituted in 200 μl DMSO, then 300 μl of base solution was added. To this, 100 μl iodomethane was added, and samples were mixed using a shaker for 15 min. 2 ml LC-MS grade water was then added to quench the reaction, followed by 2 ml dichloromethane. The solution was mixed vigorously for 30 s, then centrifuged at 3,000 × g for 2 min. The upper water layer was removed, and another 2 ml of water was added. This process was repeated four times. The lower layer was then transferred to a clean glass tube and dried under N_2_ stream. Dried samples were reconstituted in 20 μl methanol for MALDI-TOF-MS and LC-MS/MS analysis.

For MALDI analysis, 2 μl of the per-O-methylated N-glycans was mixed with 2 μl of DHB matrix (15 mg/ml in 70/30/0.1 acetonitrile / water / formic acid). The sample was then spotted on a MALDI plate and allowed to dry. Samples were analyzed using an AB Sciex TOF/TOF 5800 mass spectrometer in reflector mode. MALDI-TOF-MS data was analyzed using Data Explorer™ software and GlycoWorkBench.

Per-O-methylated N-glycans were mixed in 50/50 MeOH/H_2_O for LC-MS/MS analysis. Samples were analyzed on a Thermo Fisher Orbitrap Fusion Tribrid mass spectrometer equipped with a nanospray ion source and connected to a Dionex binary solvent system (Waltham). Prepacked nano-LC columns of 15 cm in length with 75 μm internal diameter filled with 3 μm C18 material (reverse phase) were used for chromatographic separation of the glycans. LC-MS/MS runs were conducted for 72 min using a sodiated buffer system (Buffer A: 1 mM NaOAc in H2O, Buffer B: 80% ACN, 0.1% formic acid, 1 mM NaOAc). Precursor ion scan was acquired at 120,000 resolution in the Orbitrap analyzer, and precursors at a time frame of 3 s were selected for subsequent MS/MS fragmentation in the Orbitrap analyzer at 15,000 resolution. MS/MS fragmentation was conducted with fixed CID (Collision Energy 40 %) using a data-dependent scan (DDS) program, which performs an MS/MS acquisition for the most abundant ions in the MS^1^ spectrum. MS^3^ experiments were performed using a targeted approach, where an MS^3^ event was triggered when specific fragment ions (m/z 1084, 1209, 1614, 1802) were noted in the MS/MS spectrum. Precursors with an unknown charge state, or charge state of + 1 were excluded, and dynamic exclusion was enabled (30 s duration). LC-MS/MS data was analyzed using Thermo Fisher FreeStyle and GlycoWorkBench.

### Overexpression and purification of POFUT1 and POFUT2 substrates

Huh7 cells were seeded on 10 cm plates and grown to 70 % confluency. They were transfected with either mN1 EGF1-5 myc His×6 in pSecTag2 that encodes EGF-like repeats or hTSP1 TSR1-3 myc His×6 in pSecTag2 that encodes TSR using ViaFect transfection reagent following the manufacturer’s instructions. To transfect each plate, we used 10 μg of DNA and 20 μl of the transfection reagent.

Within 24 h after the transfection, cells were washed four times with DPBS, and the medium was replaced with 5 ml of serum free labeling medium. After 12 h medium was collected and recombinant proteins were purified on TALON metal affinity resin (Takara). 200 μl of the resin was loaded onto 1 ml column (Bio-Rad) and washed 3 times with 1 ml of 100 mM TBS buffer pH 8.5 with 0.5 M NaCl and 5 mM imidazole. Next, medium was diluted with an equal volume of 2 × equilibration buffer and loaded on the column. The resin was washed 3 times with 1 ml of equilibration buffer. Unbound material and all the washes were collected to the 15 ml tube and concentrated on 3 kDa cut-off filters as described for medium samples in the section GC-MS analysis and sample preparation. Proteins were eluted using 1 ml 100 mM TBS buffer pH 8.5 with 0.5 M NaCl and 0.3 M imidazole and concentrated on 3 kDa cut-off filters. To remove imidazole and exchange buffers, 450 μl of 10 mM HEPES pH 7.0 was run through the filter for 5 times. After the sample was collected from the filter, 10 % was loaded on the 15 % SDS-PAGE gel. To visualize purified proteins, gels were stained with 0.5 % Coomassie Brilliant Blue G250 solution in 50 % methanol and 10 % acetic acid, which was washed out with 10 % acetic acid. The remaining 90 % of the sample was lyophilized and hydrolyzed with 2 N TFA. After drying down samples were stored in −20 °C until GC-MS analysis. Acid hydrolysis, derivatization and GC-MS analysis were performed as described in the previous section.

## FIGURE LEGENDS

**Supplementary Figure 1. Analysis of mannose, glucose and fucose incorporation into N-glycan-associated fucose**. (A) GC chromatogram of derivatized fucose. (B) MS fragmentation of fucose. Upper panel, MS fragmentation of the peak with elution time 11.6 min. Lower panel, MS fragmentation of the peak with elution time 11.7 min. (C) Chemical structure of fucose derivatized with BSTFA/PFBO before MS fragmentation. (D and E) Incorporation of 5 mM ^13^C-UL-glucose, 50 μM ^13^C-3,4-mannose and ^13^C-6-fucose into fucose associated with N-glycoproteins produced by A549, CaCo2, HEK293, HCT116 and CHO-Lec13 cells expressed as a percentage of labeling (D) or as a percentage of maximal signal (E); HCT116 and CHO-Lec13 are GMDS mutants with inactive *de novo* pathway; n = 3; data are presented as mean ± SD. (F and G) Incorporation of 5 mM glucose, 50 μM mannose and fucose into fucose associated with N-glycoproteins produced by HepG2 cells expressed as a percentage of labeling (F) or as a percentage of maximal signal (G); monosaccharides with different number and position of ^13^C carbons were used to evaluate a possibility of kinetic isotope effect; n = 3; data are presented as mean ± SD. (H and I) Incorporation of 5 mM ^13^C-5,6-glucose, 50 μM ^13^C-4-mannose and ^13^C-UL-fucose into fucose associated with N-glycoproteins produced by HepG2 cells during 48 h labeling period expressed as a percentage of labeling (H) or as a percentage of maximal signal (I); n = 3; data are presented as mean ± SD.

**Supplementary Figure 2. HPLC analysis of GDP-fucose pool size.** (A) HPLC separation of nucleotide and nucleotide sugar standards. (B) Standard curve presenting linear correlation between an amount of injected GDP-fucose and an area of the peak eluting from the column. (C) HPLC profile of nucleotide sugars isolated from HepG2, Huh7 and CHO cells untreated with fucose (black) and treated with 50 μM fucose (red); green arrow indicates UDP-arabinose and red arrow indicates GDP-fucose. (D) Zoom into elution time that covers GDP-fucose only.

**Supplementary Figure 3. Analysis of mannose and glucose incorporation into N-glycan-associated fucose.** Incorporation of 5 mM ^13^C-UL-glucose and 50 μM ^13^C-3,4-mannose into mannose associated with N-glycoproteins produced by HeLa, HepG2, Huh7, CHO, A549, CaCo2, HEK293, HCT116 and CHO-Lec13 cells expressed as a percentage of maximal signal; n = 3; data are presented as mean ± SD.

**Supplementary Figure 4. Analysis of newly synthesized proteins secretion and incorporation of exogenous fucose into different positions in N-glycan.** (A) Secretion of ^35^S Met labeled, newly synthesized proteins and fucosylated glycoproteins (LCA bound) produced by HepG2 cells; n = 3; data are presented as mean ± SD. (B) Secretion of ^35^S Met labeled, newly synthesized, fucosylated glycoproteins, pulled down with LCA produced by HepG2 cells growing in a presence or absence of 5 μg/ml BFA; n = 3; data are presented as mean ± SD. (C) Secretion and intracellular utilization of ^3^H-fucose in pre-labeled HepG2 cells grown in a presence of 10 μM cold fucose; n = 3; data are presented as mean ± SD. (D) LC-MS analysis of exogenous ^13^C-UL-fucose incorporation into cell associated N-glycans produced by A549, CaCo2, HEK293 and CHO-Lec13 cells that have only one fucose residue as well as N-glycans with two fucose residues; M+6 refers to N-glycans which incorporated a single molecule of exogenous fucose; M+12 refers to N-glycans which incorporated two molecules of exogenous fucose; CHO-Lec13 cells are GMDS mutants with inactive *de novo* pathway; data are presented as mean ± SEM; N refers to the number of unique N-glycan structures.

**Supplementary Figure 5. MALDI-TOF-MS and LC-MS/MS analysis of N-glycans secreted by HepG2 cells.** (A) MALDI-TOF-MS analysis of N-glycans secreted by HepG2 cells, growing without ^13^C-UL-fucose. (B) MALDI-TOF-MS analysis of N-glycans secreted by HepG2 cells, growing in a presence of 50 μM ^13^C-UL-fucose. (C) MALDI-TOF-MS distribution of ^12^C-fucose and ^13^C-UL-fucose for one representative N-glycan. (D) LC-MS^3^ (m/z 1334, m/z 1209) analysis of a bifucosylated N-glycan from HepG2 cell growing in 20 μM ^13^C-UL-fucose, which incorporated only one ^13^C-UL-fucose fucose residue. Inset zooms in on low m/z range.

