## Supplementary figures and images for "Metabolic heritage mapping: heterogenous pools of cytoplasmic nucleotide sugars are selectively utilized by various glycosyltransferases"

### Supplementary Figure 1

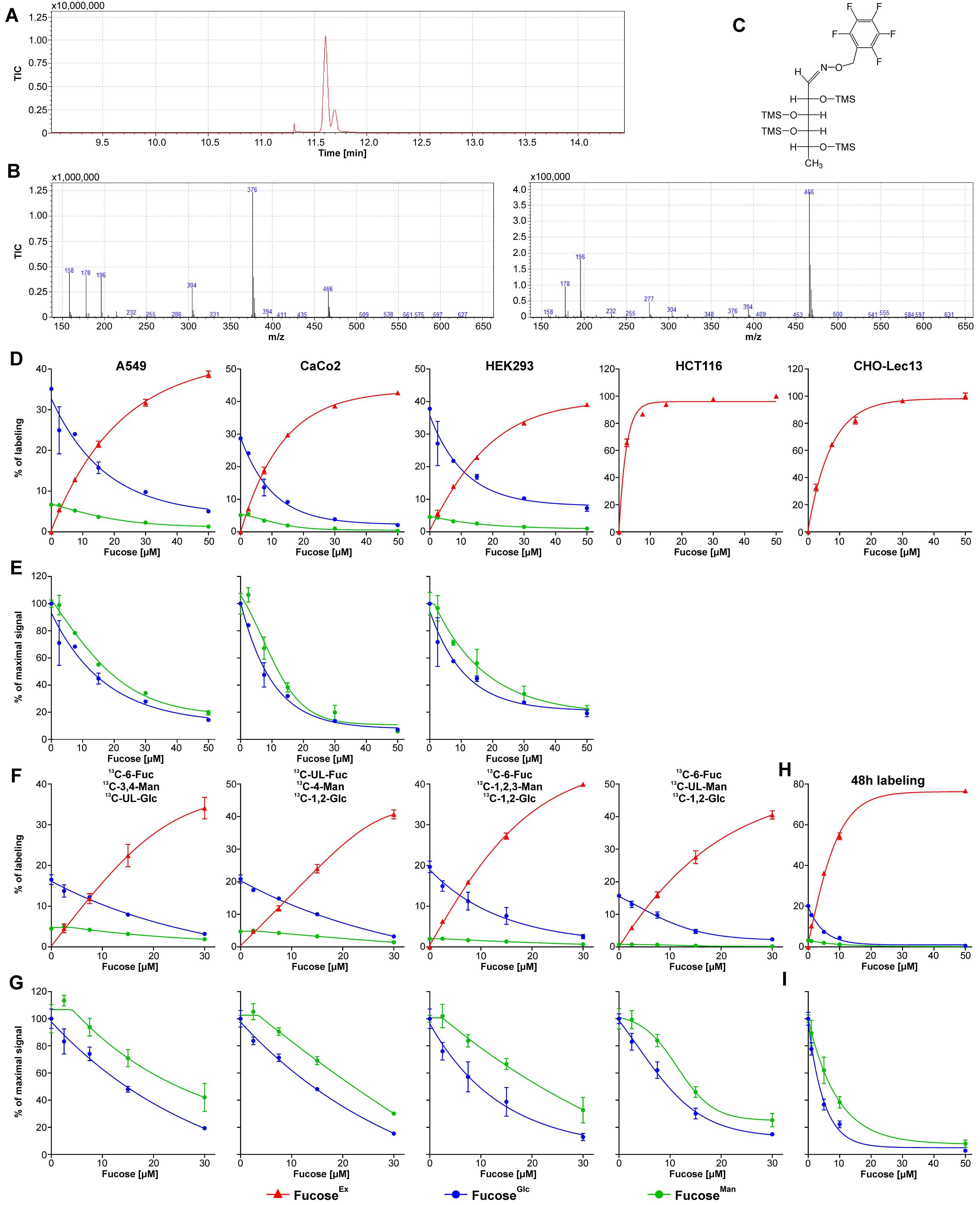

### Supplementary Figure 2

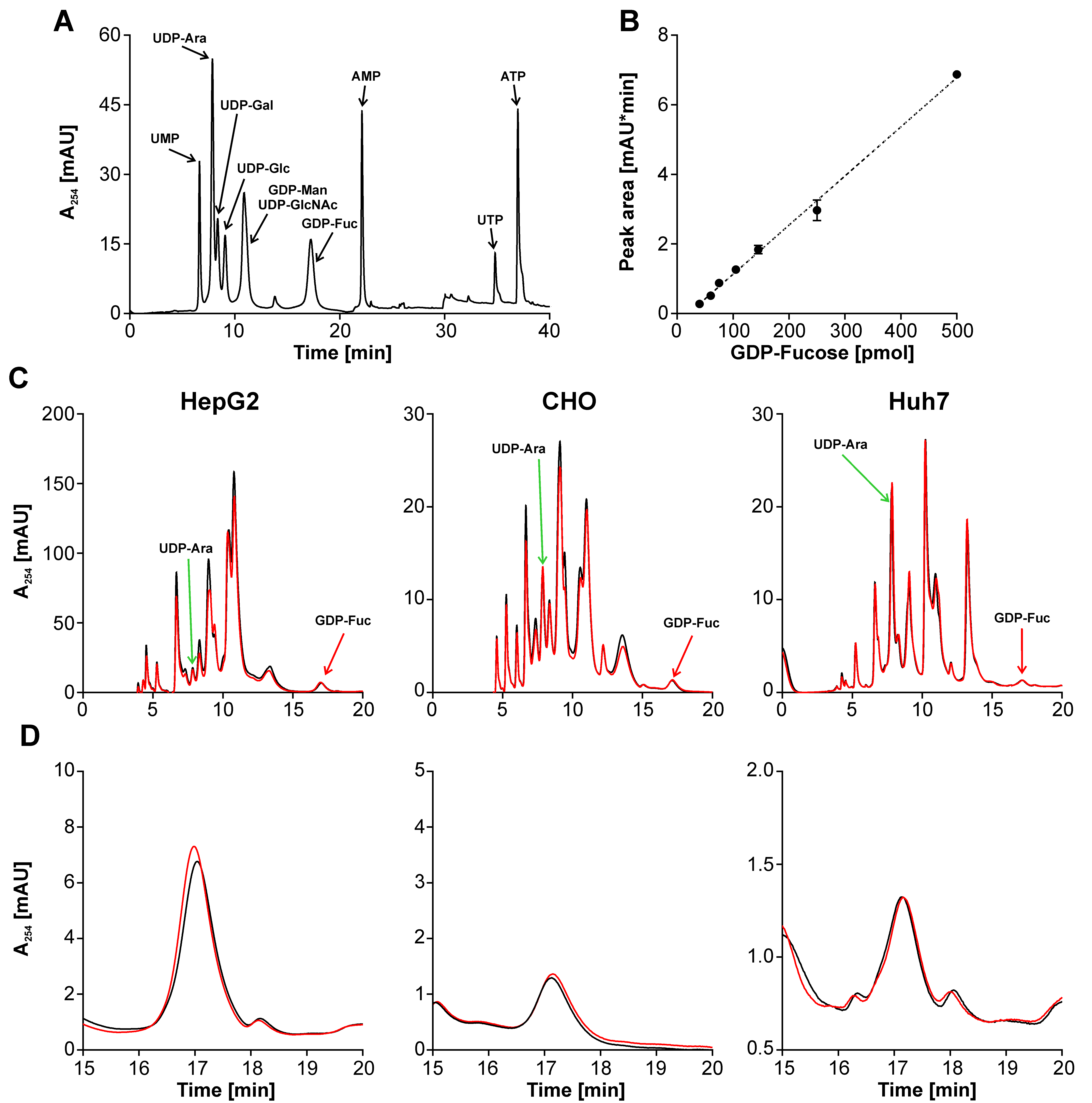

### Supplementary Figure 3

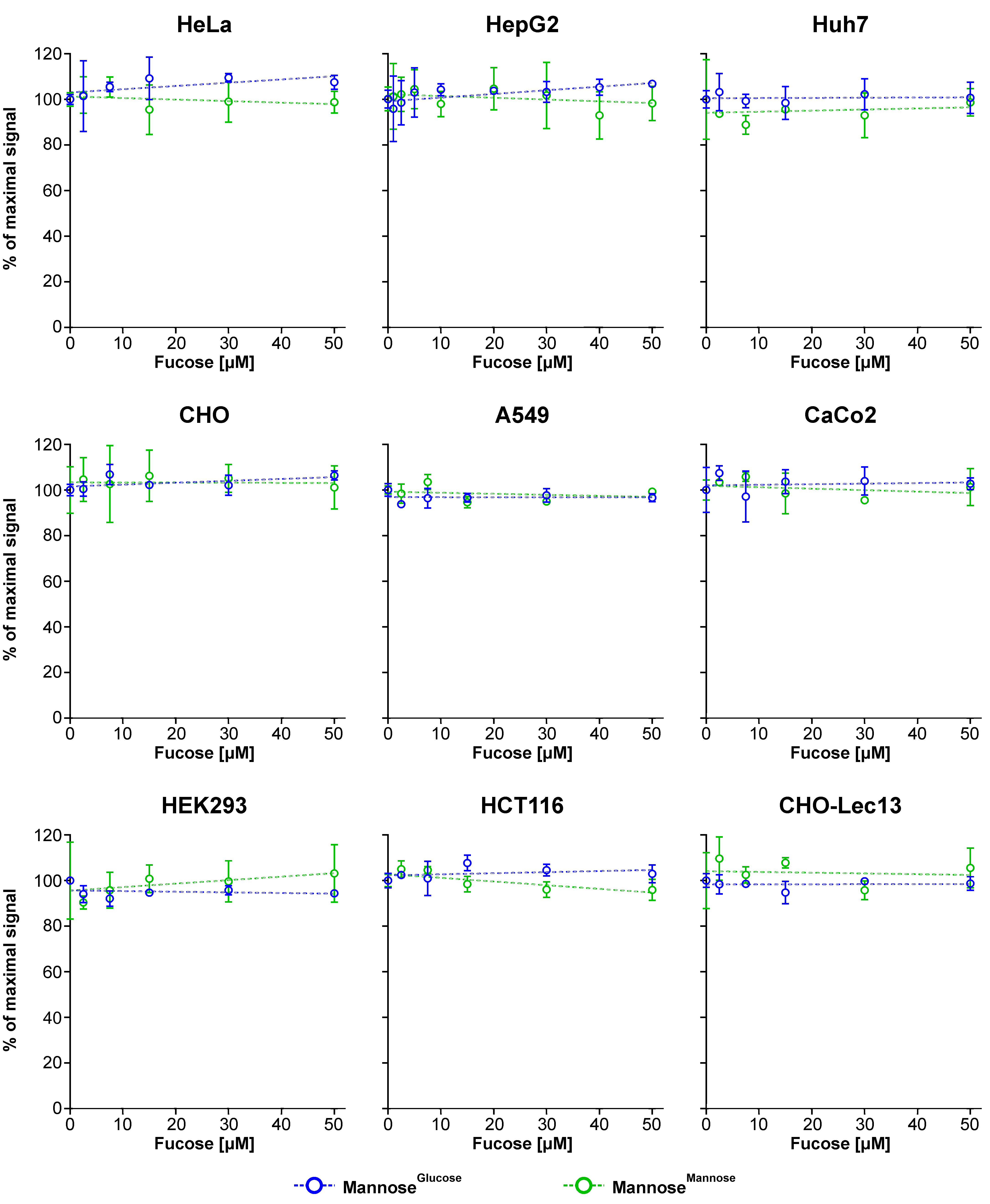

### Supplementary Figure 4

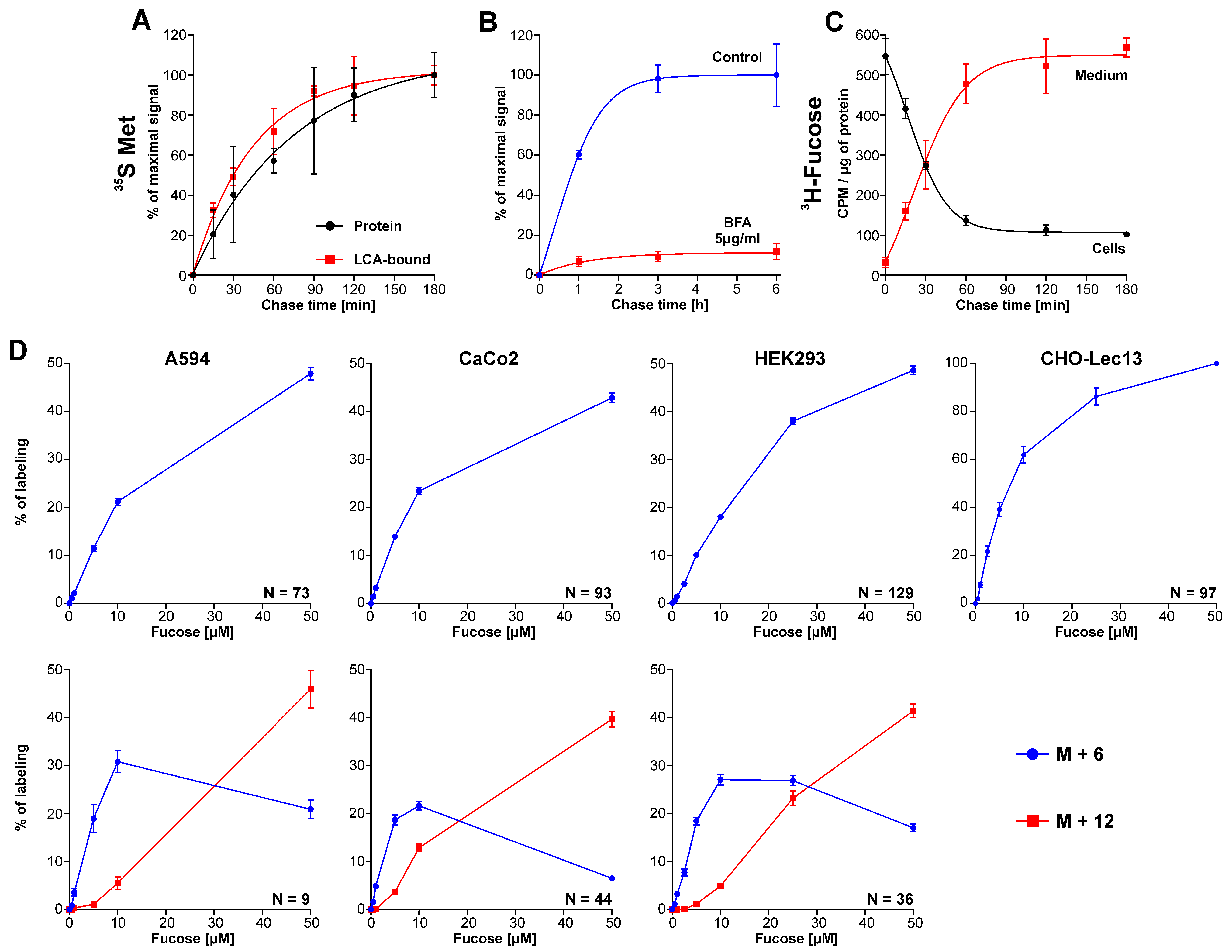

### Supplementary Figure 5

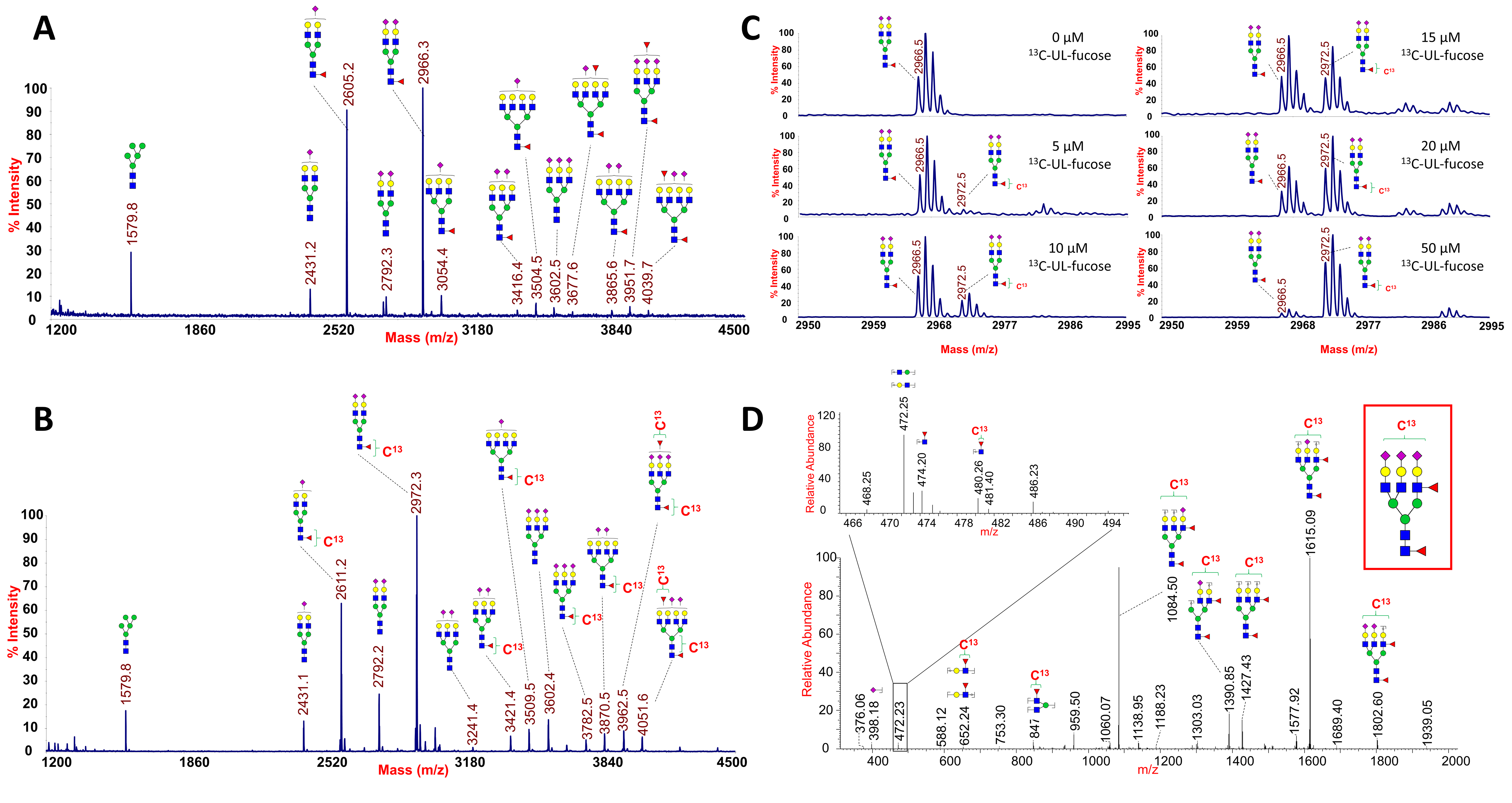
